# Long-term cognitive impairment without diffuse axonal injury following repetitive mild traumatic brain injury in rats

**DOI:** 10.1101/695718

**Authors:** Sai Ambika Tadepalli, Zsolt Kristóf Bali, Nóra Bruszt, Lili Veronika Nagy, Krisztina Amrein, Bálint Fazekas, András Büki, Endre Czeiter, István Hernádi

**Affiliations:** Department of Experimental Zoology and Neurobiology, Faculty of Sciences, University of Pécs, Ifjúság útja 6, H-7624 Pécs, Hungary; Translational Neuroscience Research Group, Szentágothai Research Centre, University of Pécs, Ifjúság útja 20, H-7624 Pécs, Hungary; Center for Neuroscience, Szentágothai Research Centre, University of Pécs, Ifjúság útja 20, H-7624 Pécs, Hungary; Grastyán Translational Research Centre, University of Pécs, Ifjúság útja 6, H-7624 Pécs, Hungary; Institute of Physiology, Medical School, University of Pécs, Szigeti út 12, H-7624 Pécs Hungary; Department of Neurosurgery, Medical School, University of Pécs, Rét u. 2, H-7623 Pécs, Hungary; Neurotrauma Research Group, Szentágothai Research Centre, University of Pécs, Ifjúság útja 20, H-7624 Pécs, Hungary; MTA-PTE Clinical Neuroscience MR Research Group, Rét u. 2, H-7623 Pécs, Hungary

**Keywords:** chronic traumatic encephalopathy, novel object recognition, water maze, memory, glial fibrillary acidic protein, amyloid precursor protein

## Abstract

Repetitive mild traumatic brain injuries (TBI) impair cognitive abilities and increase risk of neurodegenerative disorders in humans. We developed two repetitive mild TBI models in rats with different time intervals between successive weight-drop injuries, and assessed cognitive performance and biomarker profiles. Rats were subjected to repetitive Sham (no injury), single mild (mTBI), repetitive mild (rmTBI – 5 hits, 24 h apart), rapid repetitive mild (rapTBI – 5 hits, 5 min apart) and single severe (sTBI) TBI. We assessed cognitive performance 2 and 8 weeks after TBI in the novel object recognition test (NOR), and 6-7 weeks after TBI in the water maze (MWM). Acute immunohistochemical markers were checked 24 h after TBI, and blood biomarkers were measured with ELISA 8 weeks after TBI. In the NOR, both rmTBI and rapTBI showed poor performance at 2 weeks post-injury. At 8 weeks post-injury, the rmTBI group still performed worse than the Sham and mTBI groups, while the rapTBI group recovered. In the MWM, the rapTBI group performed worse than Sham and mTBI. Acute APP and RMO-14 immunohistochemistry showed axonal injury at the pontomedullary junction in the sTBI, but not in other groups. ELISA showed increased serum GFAP levels 8 weeks after sTBI, while no differences were found between the injury groups in the levels of phosphorylated-tau and S100β. Results suggest that the rmTBI protocol is the most suitable model for testing cognitive impairment after mild repetitive head injuries. The lack of common biomarkers suggests novel unknown underlying mechanisms of rmTBI.

## 1. Background

Traumatic brain injury (TBI) is a globally acknowledged health problem, and is one of the major causes of death and disability worldwide [1], [2]. It is defined as an external blunt force trauma to the head. At the cellular level, the immediate effects of TBI, called primary axotomy, are stretch and shear injuries to axons leading to disruptions of axonal cytoskeleton, followed by secondary axotomy which causes altered electrochemical functions of the damaged neurons, and ultimately leads to neuronal death [3].

Traumatic brain injury has been shown to cause cognitive decline and behavioural changes, white matter damage, and may act as a trigger for subsequent neurocognitive disorders in old age [4], [5]. Traumatic brain injury is more likely to occur in vehicular accidents, and certain sports such as football and boxing [6]. In such sports, concussive injuries are classified as mild TBI, however, they occur frequently. Once dismissed as “harmless”, concussions have now been found to cause neuropsychological consequences, which can persist up to 3 months after the injury, often called as “post-concussion syndrome” [7], [8]. It has been suggested that victims of mild TBI who suffer repeated concussion, are likely to experience increased vulnerability to cerebral damage – a phenomenon referred to as second-impact syndrome [9]. Such repetitive brain injuries also increase the risk of developing neurocognitive disorders in old age [10]. Several studies have described functional, as well as pathologic outcomes of repetitive mild TBI, such as reactive astrogliosis and axonal damage following injury [11]–[14]. While several blood and CSF biomarkers have been proposed and used in diagnosing the extent of cerebral damage following multiple concussive injuries [15], changes in serum biomarkers over time and their post-TBI predictive value are either disputable or mostly unknown. Hence, it is necessary to identify biomarkers that can be measured in the acute as well as in the chronic period, and that may be useful for selecting patients at risk of developing neurocognitive disorders and for guiding pharmacological interventions.

Most of the currently available studies have investigated acute and sub-acute effects of repetitive mild TBI [16], [17]. Unfortunately, little is known about the cumulative effect of multiple episodes of mild TBI with different inter-injury intervals and their long-term effects. Therefore, in the present study, we developed two different repetitive mild TBI models in rats with short or long time intervals between the successive injuries, and measured the cognitive performance and established biomarkers of TBI in the animals. The goal of our study was two-fold: 1) to determine the temporal window of vulnerability of the brain to a second impact, and 2) to assess the effect of repetitive mild TBI on behavioural and molecular outcomes.

## 2. Materials and Methods

### 2.1 Animals

Adult male Long Evans rats (Charles River Laboratories, Germany, aged 8-10 months at the beginning of the study) weighing 400-500 g were used. Seventy rats were used in the behavioural tasks, while additional fifteen rats were used for post-injury 24 h immunohistochemistry. Animals were pair-housed, and were kept under controlled conditions (standard 12 h light cycle from 7 a.m. to 7 p.m., with controlled temperature and humidity). Rats were maintained at 80–85% of their free feeding weight by restricting their laboratory chow supplement. Typically, they were fed with 17 g of laboratory chow per animal per day. Water was provided ad libitum. Twelve weeks prior to the behavioural testing, all rats were regularly handled for proper acclimatization to the lab environment and experimenters. During the experiments, every effort was made to minimize distress of animals. All experimental procedures were approved by the Animal Welfare Committee of the University of Pécs and the Ministry of Agriculture of the Hungarian Government. Procedures fully complied with Decree No. 40/2013. (II. 14.) of the Hungarian Government and EU Directive 2010/63/EU on the protection of animals used for scientific purposes (License no. BA02/2000-69/2017, issued: 17 Nov 2017).

### 2.2 Experimental Traumatic Brain Injury

Animals were anaesthetised with isoflurane gas. Anaesthesia was induced for 5 min with 4% isoflurane (Forane, Abbott, Hungary) in 70% N_2_ and 30% O_2_ in an induction box, and rats were maintained under anaesthesia throughout the injury and surgical procedure. Rats were ventilated with 3% isoflurane in 70% N_2_ and 30% O_2_ (Inspira ASV, Harvard Apparatus, Holliston, USA). Through the entire surgical procedure physiologic parameters (oxygen saturation (SpO_**2**_), heart rate) of the animals were monitored by a pulse oximeter (MouseOx Plus^**®**^, Starr Life Sciences Corp., Oakmont, PA), while body temperature was monitored and kept in 37°C by a Homeothermic Monitoring System (Harvard Apparatus, Holliston, MA, USA). All the monitored physiological parameters were within the normal ranges throughout all operations.

Once the anaesthesia was stabilized, the animals were exposed to an impact acceleration method of TBI initially described for rats by Foda and Marmarou [19]. A midline incision was made to expose the skull from the bregma to the lambda craniometric points. A stainless-steel disc (10 mm in diameter and 3 mm thickness) was fixed on the skull centrally between the lambda and bregma craniometric points using cyanoacrylate adhesive, in order to reduce the risk of skull fracture. The rat was placed prone on a foam bed under a 2 m high, hollow plexiglass tube with an inner diameter of 19 mm, which contained 9 cylindrical brass weights (weighing 50 g each) that were attached to each other. The total 450g weight was dropped onto the stainless disc fixed to the skull. Severity of injury was determined as the height from which the weight was dropped (Table 1). Categorization of severity was based on our pilot study where a 15cm injury was found to cause no persistent cognitive impairment, while a 150cm injury exerted substantial memory deficit. The rat was then placed back on the stereotaxic frame to remove the disc. The exposed scalp was sutured, and the rat was placed in an empty cage for recovery. Sham animals were prepared for injury in the same fashion but were not injured (Fig. 1).

**Figure 1.**
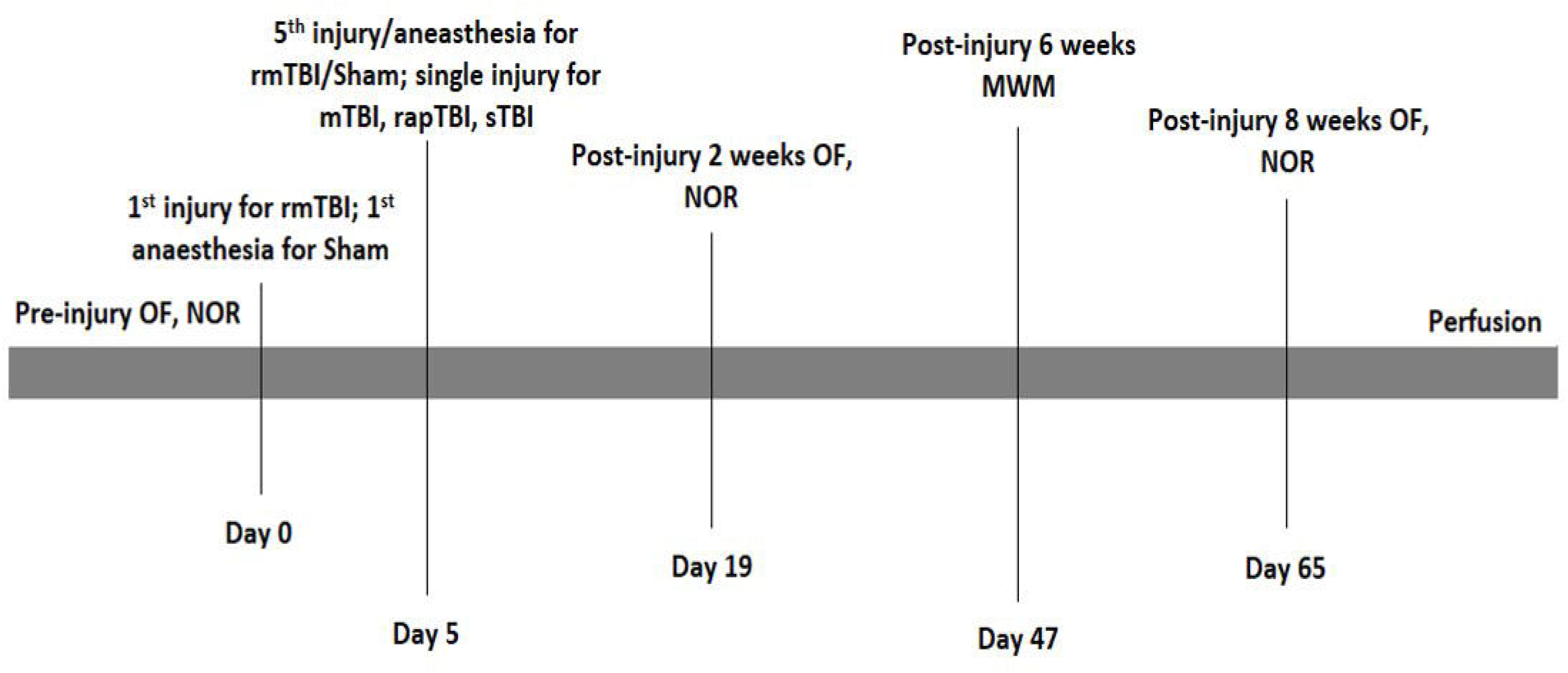
Overview of the experimental schedule.

### 2.3 Behavioural Tests

#### 2.3.1 Open Field Test

Locomotor activity was measured in the open field test (OFT) apparatus. Open field test sessions were run on the day before the novel object recognition (NOR) test sessions in order to habituate rats to the arena. The OFT was performed in an open field box which was made of black-coloured plywood, in size of 57.5×57.5 cm (length x width) surrounded by 39.5 cm high walls. The floor of the arena was divided with light grey painted lines to four by four equal squares. In each session, rats were allowed to explore the OFT arena for 5 min. After each rat, the box was thoroughly cleaned using 20% ethanol. Line crossings were counted manually and were considered as a measurement of locomotor activity. Each trial was recorded using a video camcorder (JVC super LoLux color video camera, JVC KENWOOD Corporation, Yokohama, Japan) positioned above the OFT arena, and the Ethovision XT10 tracking software (Noldus, Wageningen, Netherlands) was used for data acquisition. All animals were tested for baseline measurements (pre-injury) and at post-injury 2 weeks and 8 weeks (Fig. 1).

#### 2.3.2 Novel Object Recognition Test

Recognition memory performance of the animals was tested in the NOR test. The same apparatus (box) was used in the NOR test as in the OFT with the same video tracking system.

The NOR test included 2 trials – an acquisition trial followed by a retention trial after 30 min inter-trial interval (delay) [20]. In the first (acquisition) trial, the rats were let to explore 2 identical objects (f + f) placed in the arena for a total trial duration of 3 min. After a 30 min delay, a second (recognition) trial was run with one object identical to the sample and a novel object which had never been seen by the animal before (f + n). During the delay period, rats were not transferred back to the animal house but were kept in an empty cage in a dark room located next to the testing room. Behaviour of the animals in the second trial was also recorded for 3 min. Three different object-pairs were used. They were distributed randomly between animals and experimental sessions in a counterbalanced latin-square design.

In both trials, the time spent with the exploration of one or the other objects was recorded. The animal was considered to explore a given object, when he sniffed the object or put his nose close to it while facing the object. In the second trial, the time spent with the exploration of the novel (En) and the familiar (Ef) objects were compared by calculating a discrimination index (DI) using the following equation:

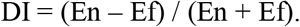

The DI was a positive number if the novel object was observed for a longer duration, while the DI was negative if the familiar object was observed for longer, and the DI was around zero if the two objects were observed for equally long time. Rats with low exploratory drive in the second trial (i.e., did not observe the two objects together for at least 5 s), or with +1.00 or −1.00 DI were excluded from the analysis.

#### 2.3.3 Morris Water Maze Test

Long-term spatial memory of the rats was tested in the Morris water maze (MWM) using a blue, circular pool, 180 cm in diameter and 90 cm in height (Ugo Basile, Gemonio, Italy). Four points around the circumference of the pool were arbitrarily designated as North, South, East, and West. On this basis, the floor area of the pool was divided into four virtual quadrants (NW, SW, SE, NE). The maze was filled with water up to the height of 30 cm, and the water was made opaque by mixing 200 g milk-powder and 30 ml blue food colouring (E131) in it. The rats were trained in the water maze task in four daily training sessions on consecutive days with four trials on each day. On training days, a hidden platform was placed in the centre of the SW quadrant. In each trial, rats were put in the water, and were allowed to search for the hidden platform for 120 s. The time elapsed until finding the platform (i.e., sitting on it) was measured as escape latency. If the platform was not found, rats were transferred to the platform at the cut-off time (120 s) and 120 s was recorded as escape latency. The quadrant from where the animal started swimming was changed clockwise in the four consecutive trials on a given day. On the fifth day, the platform was removed from the pool. A single probe trial was performed, and rats were allowed to explore the pool for 120 s. The time spent in the target quadrant was measured during the probe trial as a readout of long-term memory. Experiments were recorded using a Basler GenI acA1300 GigE camera (Basler AG, Ahrensburg, Germany). Data was processed in a PC computer, where Ethovision X10 software (Noldus, Wageningen, Netherlands) was used for recording and data analysis. Swimming behaviour of rats was automatically tracked in the Ethovision software to measure the swimming path length until finding the platform. The rats were tested in the water maze task at post-injury 6-7 weeks (Fig. 1). The overall task was divided into two weeks due to the large sample size.

### 2.4 ELISA test

Eight weeks post-injury, blood samples were obtained from all of the rats through cardiac puncture. Samples were drawn into 10ml serum separator tubes and centrifuged at 4000 rpm (1500g) for 15 min after collection. The serum was then stored at −80 °C until analysis. Commercially available sandwich ELISA kits (Elabscience®, USA) were used to measure concentration of serum pTau protein (cat. no. E-EL-R1090), GFAP (cat. no. E-EL-R1428) and S100β protein (cat. no. E-EL-R0868). 100 µl of serum samples were added to each well on the ELISA plate, and allowed to incubate for 90 minutes at 37 ºC, followed by incubation with 100 µl of biotinylated detection antibody. The plates were then washed three times with buffer and 100 µl of horseradish peroxidase-conjugate was added, followed by incubation for 30 minutes at room temperature. Finally, plates were washed three times with buffer and developed with 90 µl of substrate reagent for 15 minutes. The reaction was stopped with 50 µl of stop solution and samples were read at 450 nm with a multimode, high-performance CLARIOStar microplate reader (BMG Labtech GmbH, Ortenberg, Germany).

### 2.5 Immunohistochemistry

Twenty-four hours after the last injury (or last sham operation), 3 rats from each experimental group were euthanized with an overdose of sodium pentobarbital and were transcardially perfused with 4% paraformaldehyde containing fixative solution. On the next day, brains were removed and immersed in the same fixative overnight (16–18 h). A midline, 5mm-wide block of the brainstem was removed using a sagittal brain blocking device (Acrylic Brain Matrix for Rat, World Precision Instruments, Sarasota, FL) to include the region extending from the interpeduncular fossa to the second cervical segment. All blocks were sectioned sagittally with a Vibratome Series 1500 Tissue Sectioning System (Technical Products International Inc., St. Louis, MO) at a thickness of 40 µm and collected in PBS. Sections were collected in a serial fashion then processed for immunohistochemical localization of damaged axonal profiles via the detection of the amyloid precursor protein (APP) or neurofilament medium chain (RMO-14).

Sections were washed three times for 10 min with PBS, then treated with 0.3% hydrogen peroxide (H_2_O_2_) in PBS for 30 minutes to suppress endogenous peroxidase activity followed by washing three times in PBS. The sections were then exposed to controlled-temperature microwave antigen-retrieval in citrate buffer (pH 6.0, 0.1M) with PELCo BioWave 34700-230 (Ted Pella Inc., Redding, CA, USA). After three quick rinses in PBS, sections were immersed for 60 min in 10% bovine serum albumin (BSA) diluted in PBS containing 0.2% Triton X-100. Then, the sections were incubated overnight at 4 °C in rabbit anti-APP antibody (cat. no. 51-2700, Invitrogen, Thermo Fisher Scientific, Waltham, MA) or mouse anti-RMO-14 antibody (cat. no. 34-1000, Invitrogen, Thermo Fisher Scientific, Waltham, MA) diluted in 1% BSA/PBS to 1:1000 (APP) or 1:2000 (RMO-14) and then were washed with PBS three times for 10 min. Thereafter, the sections were subjected to the staining protocol of the Vectastain Universal Elite ABC Kit (PK-6200, Vector Laboratories, Inc., Burlingame, CA). Finally, the end product of the immunohistochemical reaction was visualized with diaminobenzidine (DAB, VWR International, Radnor, PA): sections were rinsed for 5 minutes in a 0.67 g/l DAB and 0.3 g/l H_2_O_2_ containing PBS solution. After subsequent washing in PBS 2 times for 10 min, the sections were mounted and cleared for routine light microscopic examination.

### 2.6 Statistical Analysis

All quantitative data are expressed as mean ± standard error of the mean (s.e.m). Statistical analyses were performed using IBM SPSS Statistics for Windows, Version 23 (IBM Corporation, Armonk, NY) and MS Excel (Microsoft Corporation, Redmond, WA). For analysing performance in the OFT, NOR and MWM probe-trial, one-way ANOVA test was applied to compare the injury groups, followed by Tukey’s post-hoc test. In the NOR, Student’s t-test was used to analyse the preference for the novel object above the chance level (DI = 0). Morris Water Maze acquisition data were analysed by two-factor mixed-ANOVA (Within-subject factor: DAYS. Between-subject factor: INJURY). Protein concentrations in different experimental groups measured with ELISA were compared using Kruskal-Wallis non-parametric rank test and Dunn’s post-hoc test. A level of p<0.05 was considered statistically significant in all analyses.

## 3. Results

### 3.1 Traumatic brain injury has no significant effect on locomotor activity

Locomotor activity was measured by counting line crossings in the OFT apparatus at pre-injury, post-injury 2-week and post-injury 8-week time points (Fig. 1). Animals of all injury groups exhibited overall good locomotor function in the pre-injury test (Fig. 2A), with no statistical difference in performance (F(4, 63)=1.172; p=0.332). All injury groups performed similarly in both the post-injury 2 weeks (F(4,65)=0.835; p=0.508) and the post-injury 8 weeks tests (F(4, 65)=0.138; p=0.967), indicating no major impairment in locomotor function as a result of any types of TBI.

**Figure 2:**
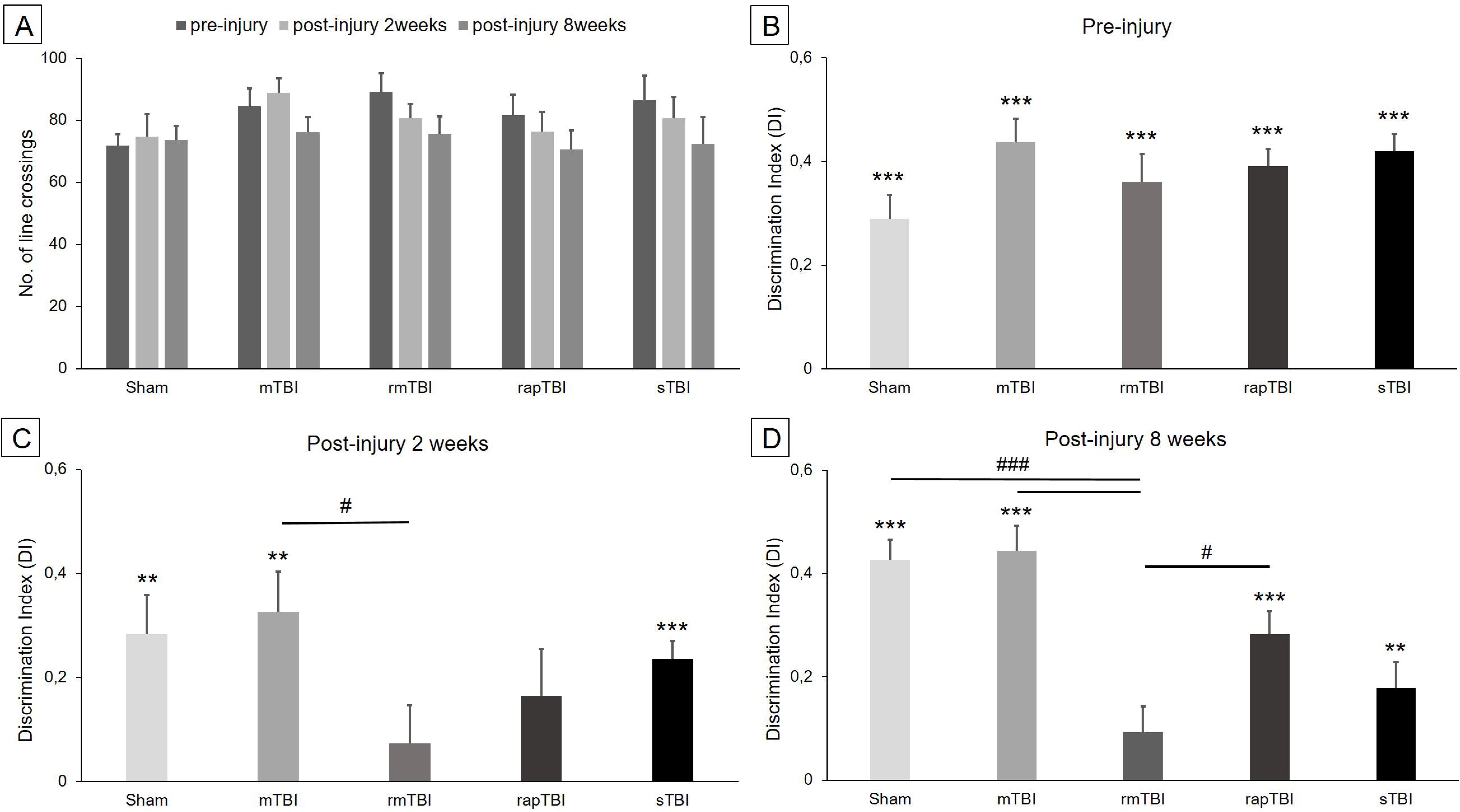
Effects of different kinds of TBI on the behavioural performance of rats in the OFT (**A**), and in the NOR task (**B-D**). A: Locomotor activity was measured by counting line crossings in the open field test (Sham and sTBI: n=13/group, mTBI, rmTBI, rapTBI: n=14/group). No gross locomotor deficits were observed in any experimental groups in any measurement points. **B**: In the pre-injury NOR test, all groups performed similarly, and were able to discriminate between familiar and novel objects (F(4, 66)=1.580; p=0.190; n=14/group). **C**: In the post-injury 2 weeks NOR test, both repetitive injury groups, rmTBI (n=11) and rapTBI (n=12), were unable to discriminate between the familiar and the novel objects, while other groups performed normally. **D**: In the post-injury 8 weeks NOR test, only rmTBI (n=12) failed to discriminate between the novel and the familiar object (p=0.09), and performed significantly worse compared to Sham (n=13; F(4, 59)=10.385; Sham vs. rmTBI: p<0.001), mTBI (n=13; mTBI vs. rmTBI: p<0.001) and rapTBI (n=13; rapTBI vs. rmTBI: p<0.05) groups.

### 3.2 Repetitive mild TBI causes irreversible long-term impairment in the NOR test

In the pre-injury NOR test, all groups were able to discriminate between the familiar and the novel objects (Fig. 2B). Discrimination index values for each group were above the chance level. For Sham: 0.289 ± 0.046 (t=6.039, df=13, p<0.001); mTBI: 0.437 ± 0.045 (t=9.226, df=14, p<0.001); rmTBI: 0.360 ± 0.053 (t=6.085, df=13, p<0.001); rapTBI: 0.391 ± 0.033 (t=10.981, df=13, p<0.001); sTBI: 0.419 ± 0.033 (t=11.225, df=13, p<0.001). All groups performed similarly in the pre-injury session (F(4, 66)=1.580; p=0.190).

In the post-injury 2 weeks NOR test (Fig. 2C), both repetitive injury groups were unable to discriminate between the familiar and the novel objects (rmTBI: 0.073 ± 0.072, t=0.896, df=10, p=0.396, and rapTBI: 0.165 ± 0.09, t=1.640, df=11, p=0.129), while other groups performed normally (Sham: 0.283 ± 0.075, t=3.456, df=10, p<0.01; mTBI: 0.326 ± 0.076, t=4.108, df=12, p<0.01; sTBI: 0.235 ± 0.034, t=6.032, df=10, p<0.001). Analysis of variance statistics (ANOVAs) did not show a main effect of TBI 2 weeks after the trauma (F(4, 53)=1.556, p=0.200).

In the post-injury 8 weeks NOR test (Fig. 2D), the rmTBI group still failed to discriminate between the novel and the familiar objects (0.092 ± 0.049, t=1.857, df=11, p=0.09), while other mild injury groups performed significantly above the chance level (mTBI: 0.444 ± 0.048, t=9.082, df=12, p<0.001; rapTBI: 0.282 ± 0.045, t=6.257, df=13, p<0.001). The rmTBI group also performed worse in comparison with the Sham, the mTBI and even the rapTBI groups (F(4, 59) = 10.385, p<0.001; Sham vs. rmTBI: p<0.001; mTBI vs. rmTBI: p<0.001; rapTBI vs. rmTBI: p<0.05). Although the sTBI group discriminated between the novel and familiar objects (0.178 ± 0.050, t=3.559, df=11, p<0.01), they performed significantly worse than the Sham and the mTBI groups (Sham vs. sTBI: p<0.01; mTBI vs. sTBI: p<0.01). Results indicate that the rmTBI group suffered from significant deficits in memory retention and recall, compared to the Sham and the mTBI groups, while the rapTBI group recovered.

### 3.3 Rapid repetitive mild TBI causes deficits in recall of spatial learning in the MWM

For the acquisition phase data, results of mixed-ANOVA for escape latency indicated that there was no interaction effect between the injury groups and the training days (F(12, 192)=0.610; p=0.832) (Fig. 3A). Also, tracking of swimming path length did not show any interaction between the injury groups and the training days (F(12, 195)=0.470; p=0.931), suggesting that non of the injuries altered the swimming strategy of the animals (Fig. 3B). However, there was a significant decrease of escape latency in all groups during the training days (F(3, 192)=29.668; p<0.05), suggesting that rats in all injury groups took less time to find the platform on day 4, compared to day 1. Assessment of reference memory in the MWM probe trial was measured in terms of time spent in the target quadrant during the probe trial on day 5 (Fig. 3C). Compared to Sham and single mTBI groups, only rapTBI group performed significantly worse (F(4, 65)=4.111; p<0.01; Sham vs. rapTBI: 45.273 s ± 2.261 vs. 34.516 s ± 1.907, p<0.05; mTBI vs. rapTBI: 44.489 s ± 2.535 vs. 34.516 s ± 1.907, p<0.05). The single mTBI group performed similar to the Sham group, indicating no effect of single mTBI on spatial learning and memory (Sham vs. mTBI: 45.273 s ± 2.261 vs. 44.489 s ± 2.535, p=0.99). Surprisingly, the sTBI group did not perform worse than the Sham group (Sham vs. sTBI: 45.273 s ± 2.261 vs. 37.746 s ± 2.974, p=0.176). Based on the probe trial results, only the rapTBI group suffered from deficits in the retention of long-term spatial memory in the MWM.

**Figure 3:**
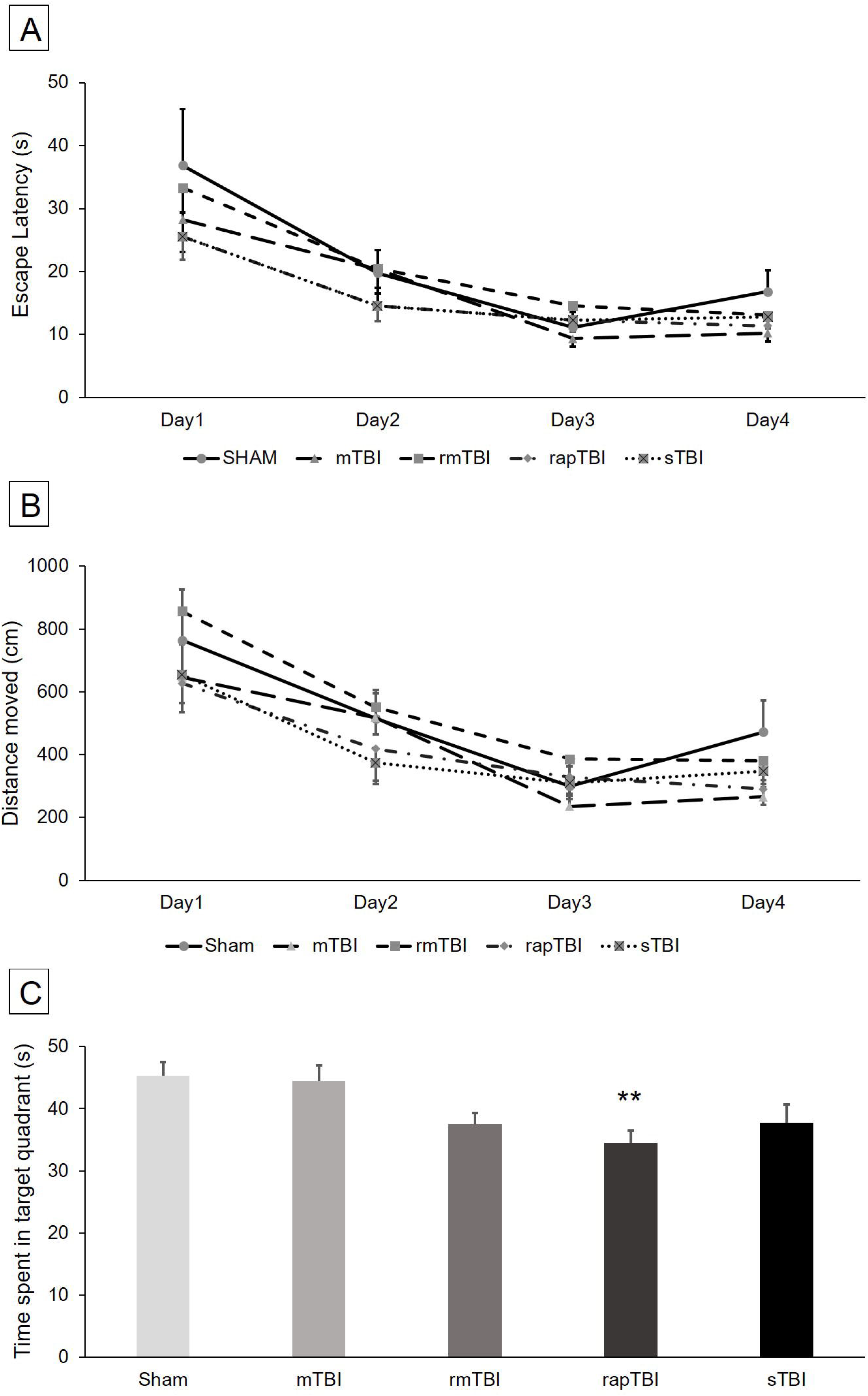
Spatial learning was tested in the Morris water maze task (MWM). Repeated measures ANOVA for escape latency (**A**) and swimming path (**B**) indicated no interaction between injury groups and experimental days (escape latency: F(12, 192)=0.610; p=0.832; swimming path: F(12, 195)=0.470; p=0.931). In the MWM probe trial (**C**), only rapTBI (n=14) performed significantly worse compared with Sham (n=13) and single mTBI (n=14) groups (F(4, 65)=4.111; p<0.01; Sham vs. rapTBI: p<0.05; mTBI vs. rapTBI: p<0.05).

### 3.4 Elevated blood GFAP levels in the severe injury group

Two months following injury, sTBI had significantly higher serum GFAP levels (Fig. 4A), compared to Sham (Kruskal-Wallis χ2=9.775, df=4, p<0.05; Sham vs. sTBI: 0.741±0.213 ng/ml vs. 2.062±0.261 ng/ml; p<0.05), mTBI (mTBI vs. sTBI: 1.115±0.259 ng/ml vs. 2.062±0.261 ng/ml; p<0.05) and rapTBI (rapTBI vs. sTBI: 1.070±0.255 ng/ml vs. 2.062±0.261 ng/ml; p<0.05). Average serum levels of GFAP in the rmTBI group were between the average levels observed in the Sham and the sTBI groups, showing a level non-significantly higher than in Sham animals but also non-significantly lower than in the sTBI group (rmTBI: 1.477±0.317 ng/ml; rmTBI vs. sTBI: p=0.135; rmTBI vs. Sham: p=0.171).

**Figure 4:**
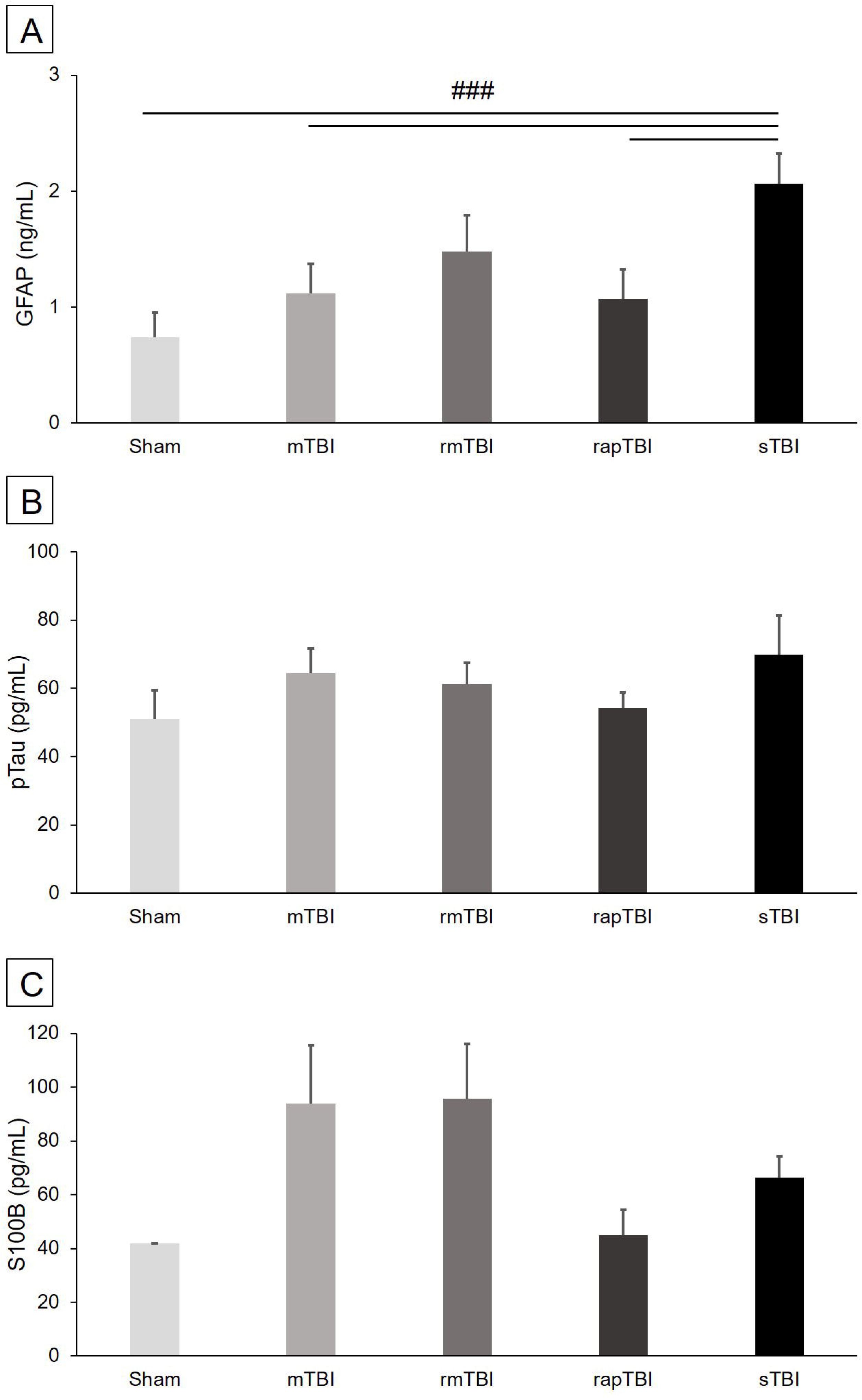
Blood biomarkers were tested 8 weeks following injury with sandwich ELISA. **A**: sTBI group (n=9) had significantly higher serum GFAP levels, compared to Sham (n=4; Kruskal-Wallis χ2=9.775, df=4, p<0.05; Sham vs. sTBI: p<0.05), mTBI (n=11; mTBI vs. sTBI: p<0.05), and rapTBI (n=11; rapTBI vs. sTBI: p<0.05) groups. **B**: No significant different was observed between injury groups in serum pTau levels (N=46; p=0.557). **C**: Similarly, no significant differences were found between injury groups in the serum levels of S100β (N=22; p=0.251).

No significant differences were observed in serum pTau levels between any injury groups (Kruskal-Wallis χ2=3.006, df=4, p=0.557) (Fig. 4B). Similarly, serum S100β levels were not found to be significantly different between all injury groups (Kruskal-Wallis χ2=5.379, df=4, p=0.251) (Fig. 4C).

### 3.5 Histological markers of axonal injury are present in sTBI but not in other TBI groups

To evaluate the axonal injury as a result of TBI of different severities, immunohistochemically labelled sections of the brainstem at the pontomedullary junction were examined under light microscope. Examination revealed few, scattered APP and RMO-14 immunopositive profiles only in the sTBI group, while other groups did not show any immunopositivity. Since only the sTBI injury group exhibited sparse APP positive and RMO-14 positive profiles, the histological markers were not further quantified. Representative images of immunohistochemical examinations are shown on Fig. 5.

**Figure 5:**
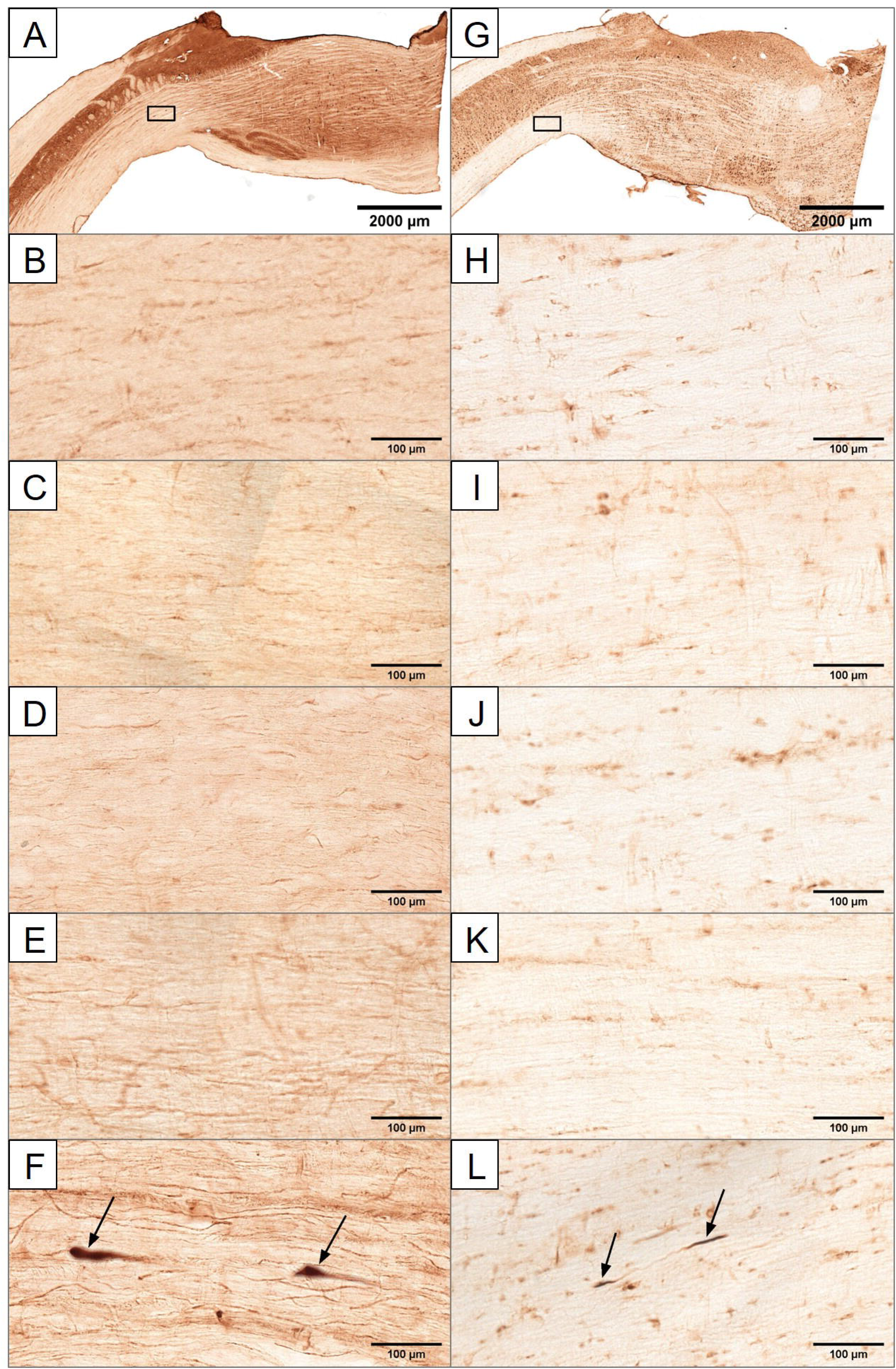
Representative photomicrographs of sagittal sections of the brainstem at the pontomedullary junction (shown on panel **A** and **G** in a large field of view) with APP (**A-F**) and RMO-14 (**G-L**) immunohistochemistry. Animals were sacrificed 24 h after TBI. Immunoreactive axonal profiles (indicated with arrows) were observed only in the sTBI group (APP: **A,F**; RMO-14: **G,L**), while Sham (APP: **B**; RMO-14: **H**), mTBI (APP: **C**; RMO-14: **I**), rmTBI (APP: **D**; RMO-14: **J**), and rapTBI groups (APP: **E**; RMO-14: **K**) did not show any APP or RMO-14 profiles. Black boxes on panel **A** and **G** shows the part from where the image on panel **F** and **L**, respectively, was captured with higher magnification.

## 4. Discussion

The current study characterizes two different models of repetitive mild TBI that replicates key functional and histological features of clinical injury in humans. To the best of our knowledge this is the first study to demonstrate in the impact acceleration model of Marmarou that a repeated mild TBI evoked from a height of 15 cm should lead to long-lasting and irreversible neurocognitive alterations. Somewhat to our surprise neurocognitive alterations were not accompanied by significant increase of the number of APP or RMO-14 immunoreactive axonal profiles at 24 h post-injury, indicating that axonal injury may not be a major player in the observed neurocognitive impairments or, alternatively, they participate in the pathology via other mechanisms. However, 8 weeks after sTBI, the glial marker GFAP that is primarily considered an acute indicator of TBI was still somewhat associated with neurocognitive impairments implicating ongoing/late onset glial pathology to the observed changes. Not surprisingly, the acute glial/BBB marker S100β did not display significant alterations at 8 weeks after TBI.

### 4.1 Behavioural alterations

In earlier studies, chronic cognitive deficits, such as depression, anxiety, and memory impairment have been observed in human TBI, as well as in experimental TBI models [7], [8], [21]. In case of a single mild TBI, memory impairment can be transient in nature. However, repetitive mild TBI could have long-lasting or irreversible effects [10]. In our study, we found acute cognitive effects in both repetitive injury groups, while Sham and single mTBI groups did not show any deficits in intermediate-term object recognition memory in the NOR test or in the spatial long-term memory in the MWM test. However, object recognition memory deficits were the most explicit in the rmTBI group, compared to other groups at the post-injury 8 weeks NOR test, while the rapTBI group recovered by that time. In contrast, long-term spatial memory deficits in the MWM probe-trial were more prominent in the rapTBI group at post-injury 6-7 weeks than in the rmTBI group.

These findings suggest that repetitive mild TBI has a chronic effect on cognitive functions regardless of the time interval between successive injuries, and mimics the functional deficits seen in humans with multiple concussive episodes. On the other hand, cognitive symptoms differ in terms of severity and the affected memory domains clearly depended on the time interval between the repetitive TBI events. Note that the OFT did not reveal any effects on basic locomotor function following injury in any of the TBI groups. Thus, we can conclude that the measures of cognitive performance were not confounded by any non-specific motor symptoms caused by TBI.

Earlier studies of repetitive mild TBI have reported behavioural deficits as well as neuropathological alterations. However, in earlier studies, much higher impact intensities were applied and were still termed as mild TBI compared to our present experiments. A 3-hit repetitive mild TBI in juvenile rats has been shown to cause short-term effects including increased axonal injury, and memory impairment in the NOR task at post-injury 28 days [22]. Repetitive mild TBI also cause cognitive impairment in mice as it was shown in the MWM test 2 months after TBI [23]. Few studies have reported chronic deficits following experimental repetitive mild TBI [12], [24], [25], lasting up to 12 months after injury in a mouse model [26]. Similar to these studies, in our study, we confirmed that repetitive mild TBI results in long-term cognitive impairment, even when the injury was caused by a very small impact. In our novel experimental design, we developed two repetitive mild TBI models, with different inter-injury intervals, to compare the progression and duration of the behavioural and pathological effects. We found that repetitive mild TBI, with a 24h inter-injury intervals (rmTBI), caused persistent cognitive deficits with no long-term histopathological changes.

### 4.2 Molecular and structural changes

Based on the ELISA results, serum GFAP levels were significantly higher in the sTBI group, compared to the Sham, mTBI and rapTBI groups. Increased expression of GFAP is a marker of astrocyte activation [27], [28]. Glial fibrillary acidic protein plays a critical role in inhibiting inflammatory response after injury effectively limiting neuronal damage [29], [30], and is a well-known acute biomarker of TBI. Previously, in a rat model of repetitive mild TBI, GFAP - in the form of reactive gliosis - was found in the cortex on the injured side 3 months following injury [31]. Indeed, the present study found increased level of GFAP in the blood of severely injured rats, while the blood level of GFAP in rats subjected to rmTBI was between the level of Sham and sTBI groups. This indicates that GFAP levels better correspond to the severity of the injury [32] than to the observed functional outcomes. Interestingly, elevated GFAP in sTBI did not coincide with significant memory loss in the post-injury 8 weeks NOR test. It is plausible that the increased GFAP level represented activated repair mechanisms following sTBI, while rmTBI induced much less extent of astrocyte activation even though they exhibited significant cognitive impairment.

Interestingly, pTau was not found to be significantly higher in any of the injured groups compared to Sham 8 weeks after TBI. Phosphorylated-tau protein, which is already well characterised in Alzheimer’s disease and in other tauopathies [33], [34], and recently has also been implicated in the pathology of TBI, where elevated levels of pTau are recognized as acute and chronic TBI biomarkers [35], [36]. Formation of tau oligomers have been observed in the brain of rats 4 hours and 24 hours following TBI [37]. While most studies reported elevated pTau protein in the cortex several weeks after the injury [26], [37], [38], one study found that transgenic mice with human tau show white matter degeneration and impaired visuospatial learning after repetitive mild TBI with only transient tau pathology in the cerebral cortex [39]. While the pathobiological role of tau in repetitive mild TBI remains a subject of extensive debate, it is likely that accumulation of tau is more pronounced at the site of injury.

S100β, a calcium-binding protein found primarily in astrocytes and Schwann cells, is used as a marker of acute glial activation [30], [40]. From our present findings, S100β levels in serum at post-injury 8 weeks was almost negligible in all injury groups. S100β protein has been found to be a sensitive biomarker, and its concentration in serum and cerebrospinal fluid (CSF) immediately after TBI has been correlated with severity and outcome of the injury [40], [41], [42], [43]. This possibly explains the small serum levels of S100β at 8 weeks after TBI in our study. Thus, it is possible that S100β is no longer expressed and/or no longer passes the BBB several weeks after TBI even if functional deficits are still present.

Twenty-four hours after the sTBI treatment, APP and RMO-14 immunoreactive profiles were observed in the white matter at the level of the pontomedullary junction of rats, indicating axonal injury. Compared to the sTBI group, no other injury groups showed explicit immunoreactive profiles, indicating minimal or no axonal damage in those groups (Sham, mTBI, rapTBI, rmTBI). Amyloid precursor protein is a well-known acute biomarker of impaired axonal transport [44], while RMO-14 is a biomarker of neurofilament compaction [45]. Mechanoporation of the axolemma – evoked by the initial shearing and tearing forces of the trauma – indicates Ca^2+^ influx into the axoplasma which may have led to impaired axonal transport and neurofilament compaction [44], [46], [47]. Both APP and RMO-14 serve as critical biomarkers for studying diffuse axonal injury (DAI) associated with TBI. Even in the medicolegal practice APP immunohistochemistry still the only “gold standard” method for detection of DAI. Based on our findings, despite the long-term cognitive impairments, rats with so mild repetitive head injury did not have any signs of DAI in the brainstem, which fact on the other hand explains the lack of motor deficits in the OFT.

### 5. Conclusions

In conclusion, here we developed two novel repetitive mild impact-acceleration TBI models, with short and long inter-injury intervals. We showed that the inter-injury interval played a crucial role in determining the extent and duration of cognitive impairment following injury. The rmTBI, with 24 h inter-injury interval, displayed long-term cognitive deficits without histological consequences, and validates the temporal window of vulnerability of the brain to a second impact. Our study reaffirms that repetitive concussive injuries with longer inter-injury intervals (24 h) causes profound and persistent neurobehavioral alterations. These results are broadly consistent with the findings of previous human studies, where repeated concussions were shown to increase the risk of chronic traumatic encephalopathy, causing chronic cognitive impairments, including severe deficits in short-term memory and executive dysfunction [10], [48]. Moving forward, the present model of rmTBI is suitable to assess the efficacy of therapeutic strategies for the management of short- and also long-term consequences of TBI in preclinical research.

## Abbreviations

ANOVA: Analysis of variance
APP: Amyloid precursor protein
BBB: Blood-brain barrier
CSF: Cerebrospinal fluid
DAI: Diffuse axonal injury
ELISA: Enzyme-linked immunosorbent assay
FGAP: Glial fibrillary acidic protein
MWM: Morris water maze
NOR: Novel object recognition
OFT: Open field test
pTau: Phosphorylated-tau
RMO-14: Neurofilament medium chain
TBI: Traumatic brain injury

## Authors’ Contributions

Conceptualization: S.T., Z.B., E.C., A.B. and I.H.; Methodology: Z.B., E.C. and I.H.; Investigation: S.T., N.B., L.N., K.A., B.F.; Formal analysis: S.T. and Z.B.; Resources: A.B. and I.H.; Supervision: Z.B., E.C., A.B. and I.H.; Funding acquisition: A.B. and I.H.; Writing—original draft preparation, S.T., Z.B. and I.H.; Writing—review and editing, E.C., K.A., A.B.; Visualization, S.T. and Z.B. All authors have approved the final manuscript.

## Acknowledgments

The authors are thankful to Kitti Göntér for her assistance in conducting the behavioural tests, and to Klára Lencse for animal care. Additionally, we are thankful to József Nyirádi for the preparation of ELISA and immunohistochemistry.

## Availability of data

The datasets used and/or analysed during the current study are available from the corresponding author on reasonable request.

## References

1. Hyder, A.A.; Wunderlich, C.A.; Puvanachandra, P.; Gururaj, G.; Kobusingye, O.C. The impact of traumatic brain injuries: a global perspective. NeuroRehabilitation 2007, 22, 341–53.

2. Selassie, A.W.; Zaloshnja, E.; Langlois, J.A.; Miller, T.; Jones, P.; Steiner, C. Incidence of long-term disability following traumatic brain injury hospitalization, United States, 2003. J. Head Trauma Rehabil. 2008, 23, 123–31.

3. Ghajar, J. Traumatic brain injury. Lancet (London, England) 2000, 356, 923–9.

4. Sivanandam, T.M.; Thakur, M.K. Traumatic brain injury: a risk factor for Alzheimer’s disease. Neurosci. Biobehav. Rev. 2012, 36, 1376–81.

5. Walker, K.R.; Tesco, G. Molecular mechanisms of cognitive dysfunction following traumatic brain injury. Front. Aging Neurosci. 2013, 5, 29.

6. Langlois, J.A.; Rutland-Brown, W.; Wald, M.M. The epidemiology and impact of traumatic brain injury: a brief overview. J. Head Trauma Rehabil. 2009, 21, 375–8.

7. Baron, D.A.; Reardon, C.L.; Defranco, J.; Baron, S.H. Concussion in Sports. In Clinical Sports Psychiatry: An International Perspective; 2013; Vol. 5, pp. 89–101 ISBN 9781118404881.

8. Ling, H.; Hardy, J.; Zetterberg, H. Neurological consequences of traumatic brain injuries in sports. Mol. Cell. Neurosci. 2015, 66, 114–22.

9. Bey, T.; Ostick, B. Second impact syndrome. West. J. Emerg. Med. 2009, 10, 6–10.

10. Stern, R.A.; Riley, D.O.; Daneshvar, D.H.; Nowinski, C.J.; Cantu, R.C.; McKee, A.C. Long-term consequences of repetitive brain trauma: chronic traumatic encephalopathy. PM R 2011, 3, S460–7.

11. DeFord, S.M.; Wilson, M.S.; Rice, A.C.; Clausen, T.; Rice, L.K.; Barabnova, A.; Bullock, R.; Hamm, R.J. Repeated mild brain injuries result in cognitive impairment in B6C3F1 mice. J. Neurotrauma 2002, 19, 427–38.

12. Ojo, J.O.; Mouzon, B.; Algamal, M.; Leary, P.; Lynch, C.; Abdullah, L.; Evans, J.; Mullan, M.; Bachmeier, C.; Stewart, W.; et al. Chronic repetitive mild traumatic brain injury results in reduced cerebral blood flow, axonal injury, gliosis, and increased T-Tau and Tau oligomers. J. Neuropathol. Exp. Neurol. 2016, 75, 636–655.

13. Uryu, K.; Laurer, H.; McIntosh, T.; Praticò, D.; Martinez, D.; Leight, S.; Lee, V.M.-Y.; Trojanowski, J.Q. Repetitive mild brain trauma accelerates Aβ deposition, lipid peroxidation, and cognitive impairment in a transgenic mouse model of Alzheimer amyloidosis. J. Neurosci. 2002, 22, 446–454.

14. Bolton Hall, A.N.; Joseph, B.; Brelsfoard, J.M.; Saatman, K.E. Repeated Closed Head Injury in Mice Results in Sustained Motor and Memory Deficits and Chronic Cellular Changes. PLoS One 2016, 11, e0159442.

15. Diaz-Arrastia, R.; Wang, K.K.W.W.; Papa, L.; Sorani, M.D.; Yue, J.K.; Puccio, A.M.; McMahon, P.J.; Inoue, T.; Yuh, E.L.; Lingsma, H.F.; et al. Acute biomarkers of traumatic brain injury: relationship between plasma levels of ubiquitin C-terminal hydrolase-L1 and glial fibrillary acidic protein. J. Neurotrauma 2014, 31, 19–25.

16. Laurer, H.L.; Bareyre, F.M.; Lee, V.M.; Trojanowski, J.Q.; Longhi, L.; Hoover, R.; Saatman, K.E.; Raghupathi, R.; Hoshino, S.; Grady, M.S.; et al. Mild head injury increasing the brain’s vulnerability to a second concussive impact. J. Neurosurg. 2001, 95, 859–70.

17. Creeley, C.E.; Wozniak, D.F.; Bayly, P. V; Olney, J.W.; Lewis, L.M. Multiple episodes of mild traumatic brain injury result in impaired cognitive performance in mice. Acad. Emerg. Med. 2004, 11, 809–19.

18. Foda, M.A.; Marmarou, A. A new model of diffuse brain injury in rats. Part II: Morphological characterization. J. Neurosurg. 1994, 80, 301–13.

19. Ennaceur, A.; Delacour, J. A new one-trial test for neurobiological studies of memory in rats. 1: Behavioral data. Behav. Brain Res. 1988, 31, 47–59.

20. Washington, P.M.; Forcelli, P.A.; Wilkins, T.; Zapple, D.N.; Parsadanian, M.; Burns, M.P. The effect of injury severity on behavior: a phenotypic study of cognitive and emotional deficits after mild, moderate, and severe controlled cortical impact injury in mice. J. Neurotrauma 2012, 29, 2283–96.

21. Aungst, S.L.; Kabadi, S. V.; Thompson, S.M.; Stoica, B.A.; Faden, A.I. Repeated mild traumatic brain injury causes chronic neuroinflammation, changes in hippocampal synaptic plasticity, and associated cognitive deficits. J. Cereb. Blood Flow Metab. 2014, 34, 1223–32.

22. Mannix, R.; Meehan, W.P.; Mandeville, J.; Grant, P.E.; Gray, T.; Berglass, J.; Zhang, J.; Bryant, J.; Rezaie, S.; Chung, J.Y.; et al. Clinical correlates in an experimental model of repetitive mild brain injury. Ann. Neurol. 2013, 74, 65–75.

23. McAteer, K.M.; Corrigan, F.; Thornton, E.; Turner, R.J.; Vink, R. Short and Long Term Behavioral and Pathological Changes in a Novel Rodent Model of Repetitive Mild Traumatic Brain Injury. PLoS One 2016, 11, e0160220.

24. Petraglia, A.L.; Plog, B.A.; Dayawansa, S.; Chen, M.; Dashnaw, M.L.; Czerniecka, K.; Walker, C.T.; Viterise, T.; Hyrien, O.; Iliff, J.J.; et al. The spectrum of neurobehavioral sequelae after repetitive mild traumatic brain injury: a novel mouse model of chronic traumatic encephalopathy. J. Neurotrauma 2014, 31, 1211–24.

25. Rubenstein, R.; Sharma, D.R.; Chang, B.; Oumata, N.; Cam, M.; Vaucelle, L.; Lindberg, M.F.; Chiu, A.; Wisniewski, T.; Wang, K.K.W.; et al. Novel Mouse Tauopathy Model for Repetitive Mild Traumatic Brain Injury: Evaluation of Long-Term Effects on Cognition and Biomarker Levels After Therapeutic Inhibition of Tau Phosphorylation. Front. Neurol. 2019, 10, 124.

26. Brenner, M. Role of GFAP in CNS injuries. Neurosci. Lett. 2014, 565, 7–13.

27. Eng, L.F.; Ghirnikar, R.S. GFAP and astrogliosis. Brain Pathol. 1994, 4, 229–37.

28. Lei, J.; Gao, G.; Feng, J.; Jin, Y.; Wang, C.; Mao, Q.; Jiang, J. Glial fibrillary acidic protein as a biomarker in severe traumatic brain injury patients: a prospective cohort study. Crit. Care 2015, 19, 362.

29. Vos, P.E.; Jacobs, B.; Andriessen, T.M.J.C.; Lamers, K.J.B.; Borm, G.F.; Beems, T.; Edwards, M.; Rosmalen, C.F.; Vissers, J.L.M. GFAP and S100B are biomarkers of traumatic brain injury: an observational cohort study. Neurology 2010, 75, 1786–93.

30. Brooks, D.M.; Patel, S.A.; Wohlgehagen, E.D.; Semmens, E.O.; Pearce, A.; Sorich, E.A.; Rau, T.F. Multiple mild traumatic brain injury in the rat produces persistent pathological alterations in the brain. Exp. Neurol. 2017, 297, 62–72.

31. Nylén, K.; Csajbok, L.Z.; Ost, M.; Rashid, A.; Blennow, K.; Nellgård, B.; Rosengren, L. Serum glial fibrillary acidic protein is related to focal brain injury and outcome after aneurysmal subarachnoid hemorrhage. Stroke 2007, 38, 1489–94.

32. Buée, L.; Bussière, T.; Buée-Scherrer, V.; Delacourte, A.; Hof, P.R. Tau protein isoforms, phosphorylation and role in neurodegenerative disorders. Brain Res. Brain Res. Rev. 2000, 33, 95–130.

33. Spillantini, M.G.; Goedert, M. Tau pathology and neurodegeneration. Lancet. Neurol. 2013, 12, 609–22.

34. Rubenstein, R.; Chang, B.; Yue, J.K.; Chiu, A.; Winkler, E.A.; Puccio, A.M.; Diaz-Arrastia, R.; Yuh, E.L.; Mukherjee, P.; Valadka, A.B.; et al. Comparing Plasma Phospho Tau, Total Tau, and Phospho Tau-Total Tau Ratio as Acute and Chronic Traumatic Brain Injury Biomarkers. JAMA Neurol. 2017, 74, 1063–1072.

35. Tsitsopoulos, P.P.; Marklund, N. Amyloid-β Peptides and Tau Protein as Biomarkers in Cerebrospinal and Interstitial Fluid Following Traumatic Brain Injury: A Review of Experimental and Clinical Studies. Front. Neurol. 2013, 4, 79.

36. Hawkins, B.E.; Krishnamurthy, S.; Castillo-Carranza, D.L.; Sengupta, U.; Prough, D.S.; Jackson, G.R.; DeWitt, D.S.; Kayed, R. Rapid accumulation of endogenous tau oligomers in a rat model of traumatic brain injury: possible link between traumatic brain injury and sporadic tauopathies. J. Biol. Chem. 2013, 288, 17042–50.

37. Cheng, J.S.; Craft, R.; Yu, G.; Ho, K.; Wang, X.; Mohan, G.; Mangnitsky, S.; Ponnusamy, R.; Mucke, L. Tau reduction diminishes spatial learning and memory deficits after mild repetitive traumatic brain injury in mice. PLoS One 2014, 9, e115765.

38. Mouzon, B.; Bachmeier, C.; Ojo, J.; Acker, C.; Ferguson, S.; Crynen, G.; Davies, P.; Mullan, M.; Stewart, W.; Crawford, F. Chronic White Matter Degeneration, but No Tau Pathology at One-Year Post-Repetitive Mild Traumatic Brain Injury in a Tau Transgenic Model. J. Neurotrauma 2018, 36, 576–588.

39. Kleindienst, A.; Hesse, F.; Bullock, M.R.; Buchfelder, M. The neurotrophic protein S100B: value as a marker of brain damage and possible therapeutic implications. Prog. Brain Res. 2007, 161, 317–25.

40. Blyth, B.J.; Farhavar, A.; Gee, C.; Hawthorn, B.; He, H.; Nayak, A.; Stöcklein, V.; Bazarian, J.J. Validation of serum markers for blood-brain barrier disruption in traumatic brain injury. J. Neurotrauma 2009, 26, 1497–1507.

41. Metting, Z.; Wilczak, N.; Rodiger, L.A.; Schaaf, J.M.; van der Naalt, J. GFAP and S100B in the acute phase of mild traumatic brain injury. Neurology 2012, 78, 1428–33.

42. Goyal, A.; Failla, M.D.; Niyonkuru, C.; Amin, K.; Fabio, A.; Berger, R.P.; Wagner, A.K. S100b as a prognostic biomarker in outcome prediction for patients with severe traumatic brain injury. J. Neurotrauma 2013, 30, 946–57.

43. Stone, J.R.; Okonkwo, D.O.; Dialo, A.O.; Rubin, D.G.; Mutlu, L.K.; Povlishock, J.T.; Helm, G.A. Impaired axonal transport and altered axolemmal permeability occur in distinct populations of damaged axons following traumatic brain injury. Exp. Neurol. 2004, 190, 59–69.

44. Marmarou, C.R.; Walker, S.A.; Davis, C.L.; Povlishock, J.T. Quantitative analysis of the relationship between intra-axonal neurofilament compaction and impaired axonal transport following diffuse traumatic brain injury. J. Neurotrauma 2005, 22, 1066–80.

45. Gaetz, M. The neurophysiology of brain injury. Clin. Neurophysiol. 2004, 115, 4–18.

46. Johnson, V.E.; Stewart, W.; Smith, D.H. Axonal pathology in traumatic brain injury. Exp. Neurol. 2013, 246, 35–43.

47. Saigal, R.; Berger, M.S. The long-term effects of repetitive mild head injuries in sports. Neurosurgery 2014, 75 Suppl 4, S149–55.

